# Michigan Neural Distinctiveness (MiND) project: Investigating the scope, causes, and consequences of age-related neural dedifferentiation

**DOI:** 10.1101/466516

**Authors:** Holly Gagnon, Molly Simmonite, Kaitlin Cassady, Jordan D. Chamberlain, Erin Freiburger, Poortata Lalwani, Shannon Kelley, Bradley Foerster, Denise C. Park, Myria Petrou, Rachael D. Seidler, Stephan F. Taylor, Daniel H. Weissman, Thad A. Polk

**Affiliations:** Department of Psychology, University of Michigan, Ann Arbor, MI, USA; Department of Radiology, University of Michigan, Ann Arbor, MI, USA; School of Behavioral and Brain Sciences, The University of Texas at Dallas, Richardson, TX, USA; Department of Applied Physiology & Kinesiology, University of Florida, Gainesville, FL, USA; Department of Psychiatry, University of Michigan, Ann Arbor, MI, USA

**Keywords:** Aging, GABA, Functional MRI, MR spectroscopy, Cognition, Individual differences, Lorazepam, Dedifferentiation

## Abstract

**Background:** Aging is often associated with behavioral impairments, but some people age more gracefully than others. Why? One factor that may play a role is individual differences in the distinctiveness of neural representations. Previous research has found that neural activation patterns in visual cortex in response to different visual stimuli are often more similar (i.e., less distinctive) in older vs. young participants, a phenomenon referred to as age-related neural dedifferentiation. Furthermore, older people whose neural representations are less distinctive tend to perform worse on a wide range of behavioral tasks. The Michigan Neural Distinctiveness (MiND) project aims to investigate the scope of neural dedifferentiation (e.g., does it also occur in auditory, motor, and somatosensory cortex?), one potential cause (age-related reductions in the inhibitory neurotransmitter gamma-aminobutyric acid (GABA)), and the behavioral consequences of neural dedifferentiation. This protocol paper describes the study rationale and methods being used in complete detail, but not the results (data collection is currently underway).

**Methods/Design:** The MiND project consists of two studies: the main study and a drug study. In the main study, we are recruiting 60 young and 100 older adults to perform behavioral tasks that measure sensory and cognitive function. They also participate in functional MRI (fMRI), MR spectroscopy (MRS), and diffusion weighted imaging (DWI) sessions, providing data on neural distinctiveness and GABA concentrations. In the drug study, we are recruiting 25 young and 25 older adults to compare neural distinctiveness, measured with fMRI, after participants take (1) a benzodiazepine (lorazepam) that should increase GABA activity or (2) a placebo.

**Discussion:** By collecting multimodal imaging measures (fMRI, MRS, DWI) along with extensive behavioral measures from the same subjects, we are linking individual differences in neurochemistry, neural representation, and behavioral performance, rather than focusing solely on group differences between young and old participants. Our findings have the potential to inform new interventions for age-related declines.

## Background

Normal aging is associated with pervasive declines in cognitive, motor, and sensory function, even in the absence of significant disease. Further, both the number of older adults and the proportion of older adults in the population are growing at alarming rates. Consequently, tens of millions of healthy people are already experiencing age-related behavioral impairments, and that number is only going to grow. Nevertheless, there are substantial individual differences in age-related behavioral impairments. Some otherwise healthy people experience significant age-related declines, while others do not. What distinguishes those who age gracefully from those who experience significant impairments? The answer to that question could transform efforts to reduce, or even reverse, behavioral impairments associated with aging.

One factor that may play an important role in explaining individual differences in aging is neural distinctiveness. Neural distinctiveness refers to the extent to which neural activation patterns evoked by different stimuli are distinguishable [1]. If two stimuli elicit activation in relatively disjoint neural populations, then the representations of those stimuli are quite distinct. Conversely, if the activated populations overlap substantially, then the representations are not very distinct. Functional neuroimaging data suggest that neural activation patterns in response to different stimuli are significantly less distinct in older compared with younger adults, a phenomenon referred to as age-related neural dedifferentiation [2–4]. Furthermore, older adults who exhibit preserved neural distinctiveness have been found to perform better than other older adults on a range of fluid processing tasks [4].

Most of the previous evidence for neural dedifferentiation has been found in the visual cortex during visual tasks. An important open question is the extent to which dedifferentiation extends to other brain regions and tasks. Single-neuron recording studies suggest that somatosensory [5,6] and auditory representations [7] become less distinct in senescent animals. While evidence in humans remains sparse, recent studies hint that age-related neural dedifferentiation occurs outside of the visual cortex and during non-visual tasks in humans as well. Payer and colleagues [8] reported age-related declines in neural distinctiveness in the ventral visual cortex as well as the prefrontal cortex during memory encoding. Neural dedifferentiation has also been observed in the inferior parietal cortex and in the medial and lateral prefrontal cortex using a whole-brain multivariate searchlight analysis [2]. Reduced distinctiveness has also been reported in the motor activity evoked by left vs. right hand tapping in older vs. younger subjects [9].

These findings suggest that age-related neural dedifferentiation may indeed be a general feature of the aging brain. The first aim of our main study is to test that hypothesis. Specifically, we are testing whether neural representations are less distinct in old than in young adults in a variety of task domains (vision, hearing, touch, motor control) and brain regions (visual cortex, auditory cortex, somatosensory cortex, motor cortex). We will also evaluate cross-domain relationships in neural distinctiveness: do older adults with less distinct visual representations also exhibit less distinct motor, somatosensory, and auditory representations? This issue has important implications for theoretical models of cognitive aging. Common-cause theories argue that age-related declines occur in tandem across domains, but process-specific theories predict that different abilities decline independently [10].

Previous work has demonstrated that neural representations become less distinctive in old age, but what causes this neural dedifferentiation? Evidence from work in non-human primates suggests that age-related reductions in the inhibitory neurotransmitter gamma-aminobutyric acid (GABA) may play a role. Leventhal and colleagues [1] demonstrated a relationship between GABA activity and neural selectivity in the visual cortex of old and young macaques. At baseline, visual neurons in older macaques responded non-selectively to orientation, showing strong responses to stimuli at a variety of different orientations. However, just minutes after the electrophoretic application of either GABA or the GABA agonist muscimol, these same cells showed strong selectivity for stimulus orientation. These effects disappeared over time, or immediately with the application of the GABA antagonist bicuculline. Conversely, visual neurons in young macaques were strongly orientation-selective at baseline, and they remained so after application of GABA. However, the application of the GABA antagonist bicuculline abolished visual selectivity in these cells and made them look like the neurons of old macaques at baseline.

Given these findings, our second aim is to investigate the relationship between GABA and neural distinctiveness in humans. In our main study, we are using magnetic resonance spectroscopy (MRS) to measure individual differences in GABA concentrations. We predict that GABA levels will be lower in older participants compared with younger participants and that participants with higher levels of GABA in specific cortical regions will exhibit greater neural distinctiveness in those same regions. In a linked drug study, we are manipulating GABA activity pharmacologically in a subset of participants to assess the impact of this manipulation on neural distinctiveness. We predict that increasing GABA activity via a low oral dose of a benzodiazepine (lorazepam, 0.5 mg) will lead to increased neural distinctiveness within individual subjects.

The third aim is to test whether individual differences in neural distinctiveness predict individual differences in behavior, particularly in older participants. Park et al. [4] found that individual differences in neural distinctiveness in the visual cortex predicted performance on a range of fluid processing tasks in older adults. In fact, neural distinctiveness accounted for 30% of the variance in behavioral performance, despite the fact that neural distinctiveness was only measured in visual cortex using simple visual tasks while the fluid processing tasks required far more general types of cognitive processing. In order to investigate the behavioral consequences of neural distinctiveness more thoroughly, we propose to collect a full battery of cognitive and sensorimotor measures in all of the participants in the main study.

## MiND Project – Main Study

The goal of the main study is to evaluate the scope of neural dedifferentiation, whether age-related declines in GABA may be a cause, and its behavioral consequences.

## Methods and design

### Participants

All participants are healthy, right-handed, native English speakers. Participants are aged 18-29 years (young adults) or 65 years and older (older adults). Major exclusion criteria are listed in Table 1. All sessions take place at the University of Michigan’s Functional MRI Laboratory at the Bonisteel Interdisciplinary Research Building and the Ann and Robert H. Lurie Biomedical Engineering Building in Ann Arbor, Michigan. Participants are being recruited from the Ann Arbor community and the surrounding area.

**Table 1.**
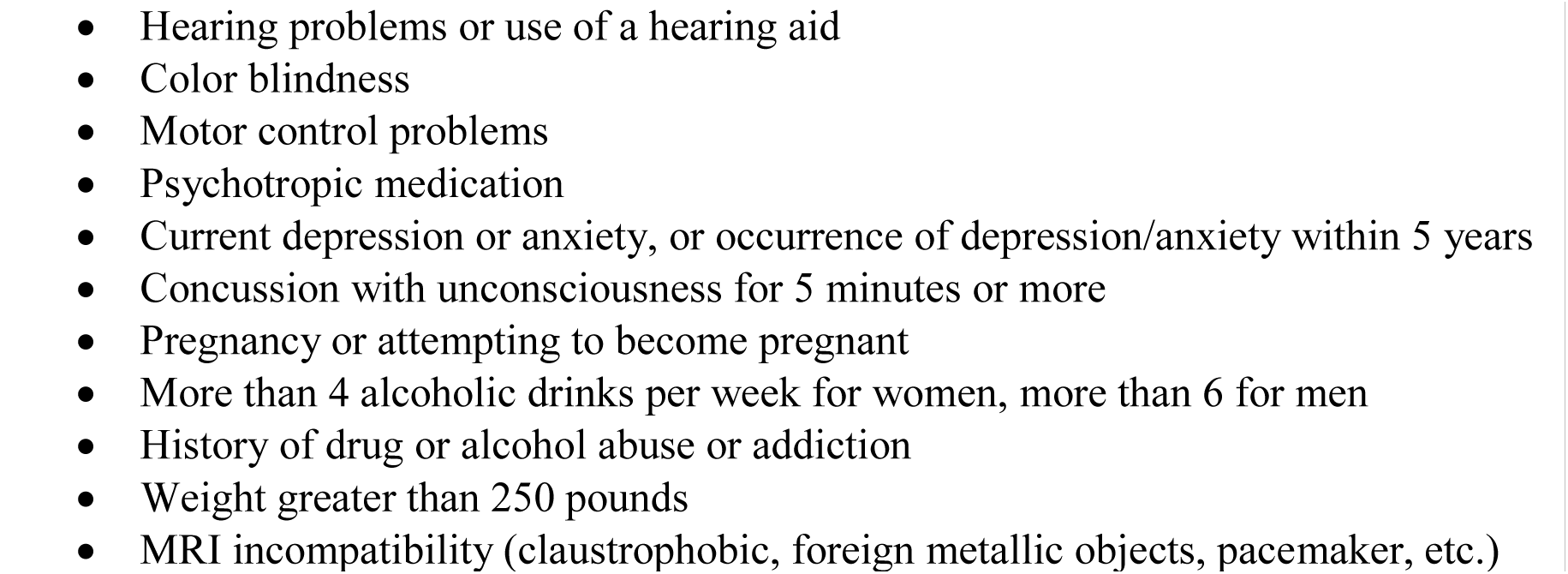
Main study exclusion criteria

|  |
| --- |
| • Hearing problems or use of a hearing aid<br>• Color blindness<br>• Motor control problems<br>• Psychotropic medication<br>• Current depression or anxiety, or occurrence of depression/anxiety within 5 years<br>• Concussion with unconsciousness for 5 minutes or more<br>• Pregnancy or attempting to become pregnant<br>• More than 4 alcoholic drinks per week for women, more than 6 for men<br>• History of drug or alcohol abuse or addiction<br>• Weight greater than 250 pounds<br>• MRI incompatibility (claustrophobic, foreign metallic objects, pacemaker, etc.) |

### Power Calculations

Carp, Park, Polk, et al. [2] found that the neural representations of visual stimuli are less distinct in older adults than in young adults (d = 1.06). To achieve 80% power to detect an effect of this size, a sample of approximately 15 subjects per group would be required. Park et al. [4] reported correlations between neural distinctiveness and fluid intelligence among older adults ranging from r = 0.275 to r = 0.59. To achieve 80% power to detect a correlation of r = 0.275, approximately 100 subjects would be required per group; to detect a correlation of r = 0.59, approximately 30 subjects would be required. Thus, to provide sufficient power to detect both between-group neural differences and brain-behavior correlations within the older group, we are targeting a sample of 100 older adults and 60 young adults.

### Session Design

After completing an initial telephone screening interview and being determined eligible, all subjects participate in three separate sessions. Session 1 lasts two hours and consists only of cognitive and behavioral tasks. Session 2 includes 45 minutes of behavioral testing and an hour-long functional magnetic resonance imaging (fMRI) scan. Session 3 includes a 1.5-hour MRS scan.

### Cognitive and Behavioral Tasks

Several tasks are being administered to assess sensory and cognitive abilities. The tasks are described below and are grouped by domain. All tasks referencing the NIH Toolbox are administered on an iPad using the NIH Toolbox^®^ for Assessment of Neurological and Behavioral Function iPad App [11]. Detailed information about the NIH Toolbox scoring methodology can be found in the scoring guide located on the NIH Toolbox website [12]. Participants complete these tasks during Sessions 1 and 2. Table 2 shows the task administration order. Participants also completed the Cognitive Failures Questionnaire during the screening process [13,14].

**Table 2.** Main study behavioral tasks and fMRI session sequence

| Session | Measure | Domain | Approx. Time (min) |
| --- | --- | --- | --- |
| 1 – Behavior Only | MoCA | Cognitive Screening Tool | 10 |
|  | VPA_1 | Episodic Memory | 9 |
|  | Faces in Noise | Visual | 2 |
|  | Auditory Threshold | Auditory Screening Tool | 7 |
|  | Sentences in Noise | Auditory | 7 |
|  | Objects in Noise | Visual | 2 |
|  | Symbol Search | Processing Speed | 4 |
|  | VPA_2 | Episodic Memory | 3 |
|  | Coding | Processing Speed | 3 |
|  | Buildings in Noise | Visual | 2 |
|  | Purdue Pegboard 1 | Motor | 5 |
|  | Digits in Noise | Auditory | 4 |
|  | Purdue Pegboard 2 | Motor | 3 |
|  | Scenes in Noise | Visual | 2 |
|  | Break |  | 5 |
|  | NIH Words-In-Noise Test | Auditory | 6 |
|  | NIH Flanker Inhibitory Control and Attention Test | Executive Function | 3 |
|  | NIH List Sorting Working Memory Test | Working Memory | 9 |
|  | NIH Dimensional Change Card Sort Test | Executive Function | 5 |
|  | NIH Pattern Comparison Processing Speed Test | Processing Speed | 3 |
|  | NIH Picture Sequence Memory Test | Episodic Memory | 6 |
|  | NIH Picture Vocabulary Test | Crystallized Intelligence | 3 |
|  | NIH Oral Reading Recognition Test | Crystallized Intelligence | 3 |
|  | NIH 9-Hole Pegboard Dexterity Test | Motor | 6 |
|  | NIH Grip Strength Test | Motor | 3 |
| 2 – Behavior + fMRI | RBMT Story 1 | Episodic Memory | 2 |
|  | Static Detection Threshold Right Hand | Tactile | 3 |
|  | Dynamic Threshold Right Hand | Tactile | 3 |
|  | Dynamic Threshold Intra-hemispheric Conditioning | Tactile | 5 |
|  | Dynamic Threshold Inter-hemispheric Conditioning | Tactile | 5 |
|  | NIH Visual Acuity Test | Visual | 2 |
|  | RBMT Story 2 | Episodic Memory | 2 |
|  | Functional Tactile Object Recognition | Tactile | 5 |
|  | NIH 2-Minute Walk Endurance Test | Motor | 6 |
|  | fMRI Task Preview |  | 10 |
|  | fMRI |  | 50 |

#### Visual Function

##### 1. NIH Toolbox Visual Acuity Test

This test measures binocular distance visual acuity. Letters appear one at a time on an iPad screen at eye level and participants view them from a distance of 3 meters. Participants verbally state the letter they see on the screen, and responses are recorded by the researcher using an iPad wireless keyboard. The letters get smaller following a correct response, and they get larger following an incorrect response. Participants are instructed to wear corrective contact lenses or glasses, if any. The software automatically calculates a LogMAR score (a modified version of a Snellen visual acuity score) and converts it to a standard score.

##### 2. Visual Tasks in Noise

Four visual tasks in noise are administered on a Dell laptop with a 15.6-inch screen using the Psychophysics Toolbox [15,16] in MATLAB [17]. The details of the four tasks are described below, but all of them consist of a fixation cross presented for 500 ms followed by a black and white picture presented in dynamic Gaussian noise for 500 ms. After the picture is presented, a response screen appears. After participants make their response, the next trial begins. The order of the stimulus presentation is pseudorandomized and is the same for each participant. Each task has 4 practice trials with feedback provided and 50 scored trials without feedback. The tasks follow a staircase procedure. When a participant makes three correct responses in a row, the next trial increases by one level of noise. Following an incorrect response, the amount of noise is decreased by one level. There are 15 levels of Gaussian noise, and each task starts at the 5^th^ level of noise. The dependent measure is the average level of noise presented for the last 40 trials.

###### a. Buildings in Noise (BIN)

The stimulus picture is either a house (50% of trials) or an apartment (50% of trials). Participants are asked to press “1” on the keyboard with their left index finger if they think the picture was a house and to press “0” with their right index finger if they think the picture was an apartment building. Building images are gathered from the same stimulus set used in Park et al. [3].

###### b. Faces in Noise (FIN)

The stimulus picture is either a male (50% of trials) or female (50% of trials) face. Participants are asked to press “1” on the keyboard with their left index finger if they think the picture was a male face and to press “0” with their right index finger if they think the picture was a female face. Face stimuli are from Gold, Bennett, and Sekuler [18].

###### c. Objects in Noise (OIN)

The stimulus picture is either an office item, such as a stapler or writing utensil (50% of trials), or a food item, such as a hamburger or salad (50% of trials). Participants are asked to press “1” on the keyboard with their left index finger if they think the picture was an office item and to press “0” with their right index finger if they think the picture was a food item. Object stimuli are from Brady, Konkle, Alvarez, and Oliva [19].

###### d. Scenes in Noise (ScIN)

The stimulus picture is either an urban (50% of trials) or nature (50% of trial) scene. Participants are asked to press “1” on the keyboard with their left index finger if they think the picture was an urban scene and to press “0” with their right index finger if they think the picture was a nature scene. Scene images are from Zhou, Lapedriza, Khosla, Oliva, and Torralba [20].

#### Auditory Function

##### 1. NIH Toolbox Words-In-Noise Test (WIN)

Measures how well participants hear words in a noisy environment. While wearing over-the-ear noise cancelling headphones, participants hear single words presented with varying levels of background noise. Words are presented separately to the right and left ears. Participants are instructed to say the word they thought they heard. The examiner indicates a correct or incorrect response on the iPad. The software automatically generates a raw score, hearing threshold, and standard score for each ear.

##### 2. Digits in Noise

This task was developed by the Nottingham University Hospitals NHS Trust and resembles the Digit Triplet Test [21]. Participants are asked to discern numbers presented in background noise. The task is administered on a Dell laptop using MATLAB [17]. Groups of 3 numbers are presented binaurally with varying signal-to-noise ratios (SNR). The level of noise is kept constant, while the decibel (dB) level of the digits varies. All participants start at 14 dB SNR. Following a correct response, the next trial is presented with the SNR decreased by 2 dB. Following an incorrect response, the next trial is presented with the SNR increased by 2 dB. Participants complete 24 trials, and their speech reception threshold is calculated by averaging the SNR dB of the last 19 trials. The task is presented using over-the-ear noise cancelling headphones, and participants adjust the volume to a comfortable level before beginning the task.

##### 3. Sentences in Noise

This task assesses how well participants can hear sentences presented in noise. The task is administered on a Dell laptop using the Oscilla USB-350SP PC-based audiometer software with TDH-39 headphones. Sentences are presented monaurally, and the volume is set to a comfortable level for each ear before starting the task. Three lists, each consisting of six sentences, are presented to each ear. After a sentence is presented, the participant is instructed to repeat back what they heard, and the researcher indicates their response on the laptop. The software automatically generates the degree of SNR loss in decibels based on the number of words the participant hears.

##### 4. Hearing Threshold

This task is administered on a Dell laptop using the Oscilla USB-350SP PC-based audiometer software with TDH-39 headphones. Hearing threshold is determined using the Hughson Westlake Automatic Hearing Test. Tones are presented monaurally at 125, 250, 500, 1000, 2000, 4000, and 8000 Hz. Participants are instructed to press a response button when they can hear a tone. The test begins by presenting a tone at 20 dB. The hearing level increases by 5 dB until the participant responds. Participants must respond to 2 out of 3 presentations of the same hearing level at each frequency before moving on to the next frequency. The minimum decibel level at which the participant responded to at least 2 out 3 presentations is recorded. The pure tone average (PTA) is calculated for each ear using the recorded decibel levels at 500, 1000, and 2000 Hz.

#### Tactile Function

##### 1. Brain Gauge Vibrotactile Tasks

A pair of piezoelectric vibrotactile stimulators (CM5, Cortical Metrics, LLC) are used to measure tactile function using the four tasks described below [22]. In all four tasks, vibrations are delivered to the pads of the index and middle fingers of the left and right hands via plastic probes measuring 5 mm in diameter. The tasks are controlled by a Windows Dell laptop using the Brain Gauge software application (Cortical Metrics, LLC). Participants use a standard computer mouse to respond to the stimuli.

###### a. Static Detection Threshold [23]

A single vibrotactile stimulation is delivered to the right index or the right middle finger at a frequency of 25 Hz. The participant indicates which of the two fingers they felt the vibration on by using the mouse with their left hand to click a button on the monitor screen. The task consists of 20 trials of stimulation; 10 stimuli are randomly presented to each finger for 500 ms. The task begins with a stimulus amplitude of 15 μm, and subsequent stimuli amplitudes are determined using a staircase procedure. For the first half of the trials, if the participant responds correctly the amplitude decreases by 1 μm; if they respond incorrectly it increases by 1 μm. For the last half of the trials, the amplitude decreases by 1 μm following two correct responses and increases by 1 μm following one incorrect response.

###### b. Dynamic Threshold [23]

Tactile stimulation is delivered to either the right index or right middle finger at a frequency of 25 Hz. Each stimulus begins at an amplitude of 0 μm then gradually increases at a rate of 2 μm per second. Once the participant can discern which finger they are feeling the vibration on, they make their response by clicking a button on the monitor screen. Participants complete a total of 7 trials, and each trial begins with a randomized delay period of 0, 1.5, 2, or 3 seconds. The stimulus amplitude at the time of the participant’s response is recorded.

###### c. Dynamic Threshold with Intra-hemispheric Conditioning

This task consists of 16 trials in which a target stimulus is delivered to the right index finger and a conditioning stimulus is concurrently delivered to the right middle finger. Similar to the dynamic threshold task previously described, the target stimulus has a starting amplitude of 0 μm and increases at a rate of 2 μm per second. The conditioning stimulus is delivered at 25 Hz with an amplitude of 15, 50, 100, or 200 μm [23,24]. There are four trials at each amplitude which are randomly presented during the task. Participants respond by pressing a computer mouse button attached to the inner right side of the device with their right thumb as soon as they are able to feel the vibration on their right index finger. The dependent variable is the stimulus amplitude at the time of the participant’s response.

###### d. Dynamic Threshold with Inter-hemispheric Conditioning

This task is the same as the intra-hemispheric task, the only difference being that the conditioning stimulus is delivered to the left index finger instead of the right middle finger. Participants press a mouse button with their right thumb as soon as they are able to feel a vibration on their right index finger.

##### 2. Functional Tactile Object Recognition [25]

In this task, participants identify everyday objects by the sense of touch. Participants place their hand in a box, preventing them from seeing the object, and the examiner places an object in their hand (soda bottle, clothespin, etc.). Participants indicate the object they think they are holding using a poster that has pictures of several different objects. Participants complete six trials. Accuracy and response time are recorded.

#### Motor Function

##### 1. NIH Toolbox 9-Hole Pegboard Dexterity Test

This is a test of manual dexterity. Participants place and remove nine pegs in a pegboard using one hand at a time. The NIH toolbox software records the time it takes to place and remove the pegs for each hand and generates a standardized score for each hand.

##### 2. NIH Toolbox Grip Strength Test

This is a measure of hand strength. Participants are seated in a chair with their feet flat on the floor. Participants squeeze a Jamar Plus Digital dynamometer as hard as they can for three seconds with their arm at a 90-degree angle. The dynamometer reports the number of pounds of force the participant generates with each hand. This measure is recorded in the software and converted to a standardized score.

##### 3. NIH Toolbox 2-Minute Walk Endurance Test

This test measures cardiovascular endurance by recording how far participants can walk in 2 minutes. Cones are placed 25 feet apart in a hallway. Participants are instructed to walk back and forth around the cones as fast as they can without running or hurting themselves for 2 minutes. The total distance walked is recorded and automatically converted to a standardized score by the software.

##### 4. Purdue Pegboard Test

Participants complete two separate tasks using the Purdue Pegboard to measure bimanual dexterity. In the “Both Hands” task, participants pick up a peg with their right hand and a peg with their left hand at the same time. They place the pegs, at the same time, in the first row of holes, and continue to place pegs down the rows. Participants are instructed to place as many pegs as they can until they are told to stop. The number of pairs of pegs placed in 30 seconds is recorded.

In the “Assembly” task, participants use both hands to create assemblies consisting of four items. Participants are instructed to 1) pick up a peg with their right hand; 2) while placing the peg in the hole with their right hand, they should pick up a washer with their left hand; 3) while placing the washer on the peg with their left hand, they should pick up a collar (a small metal tube) with their right hand; 4) while placing the collar on the peg and over the washer with their right hand, they should pick up a washer with their left hand; 4) while placing the washer on top of the collar with their left hand, completing one assembly, they should pick up a peg with their right hand to begin the next assembly. The number of items (pegs, collars, and washers) placed in 1 minute is recorded.

Each task is conducted twice, and the average number of pairs and items is recorded.

#### Cognitive Impairment Screening

##### 1. Montreal Cognitive Assessment (MoCA)

The MoCA [26] is a pen and paper based assessment tool used to assess mild cognitive impairment. It takes approximately 10 minutes to administer and consists of 13 short tasks. Visuospatial ability and executive function are assessed by a trail-making task, copying a three-dimensional cube, drawing a clock, and a verbal abstraction task. Language ability is assessed using an animal naming task, sentence repetition, and a fluency task. Memory is evaluated using a 5-word delayed recall task as well as a digits-forward and digits-backward task. Attention and concentration are measured using a target detection task and a subtraction task. Orientation is evaluated by asking the participant the date and location of the study session. Each task has a point value associated with it. The total number of points earned is recorded. The highest possible score is 30 points, and a score of 26 or higher is considered normal.

#### Processing Speed

##### 1. NIH Toolbox Pattern Comparison Processing Speed Test

Participants are instructed to discern, as fast as they can, whether two simple side-by-side pictures presented on an iPad are the same or different. Participants press buttons on the iPad screen to indicate their response. The raw score is the number of items they correctly answer in 85 seconds. The software automatically generates the raw score and converts it to a standard score.

##### 2. Wechsler Adult Intelligence Scale (WAIS-IV) - Symbol Search Subtest [27]

Each item consists of 2 target symbols adjacent to a line of 5 search symbols. Participants are instructed to draw a line through a search symbol if it matches one of the target symbols. If none of the target symbols match the search symbols, they draw a line through a “no” box. Participants complete as many items as they can in 2 minutes. Their raw score is determined by subtracting the number of incorrectly answered items from the number of correctly answered items.

##### 3. WAIS-IV – Coding Subtest [27]

A key is presented at the top of the page. In the key, each number (1-9) has its own symbol. Below the key is a grid consisting of rows of numbers. Each number has an empty space below it. The participant is instructed to draw the symbol that corresponds to each number. Participants complete as many number-symbol items as they can in 2 minutes. Their raw score is determined by subtracting the number of incorrectly answered items from the number of correctly answered items.

#### Executive Function and Working Memory

##### 1. NIH Toolbox List Sorting Working Memory Test

Pictures of different foods and animals are presented on the iPad screen one at a time. Participants are instructed to repeat the list of items in size order from smallest to largest. For the NIH scoring, in order for the participants’ response to be marked correct they must list all of the correct items in the correct order. Partial points are not awarded. The software automatically generates a raw score (the number of correct responses) and converts it to a standard score.

In order to obtain a more sensitive measure of working memory, we have devised a way to award partial points for participant responses. For each list, participants receive 1 point for each item correctly remembered regardless of order. They also receive 1 point if the first item is correct and 1 point if the last item is correct. Finally, each item is considered together with the item following it, and the participant receives 1 point if that particular pair order occurs in their response. The raw score is the total number of points they earn.

##### 2. NIH Toolbox Flanker Inhibitory Control and Attention Test

A row of arrows is presented on the iPad screen, and participants are instructed to indicate, as quickly as they can, the direction of the middle arrow. In some trials, the middle arrow points in the same direction as the arrows surrounding it (congruent trials). In other trials, the middle arrow points in the opposite direction (incongruent trials). In total, there are 20 trials, 40% of which are incongruent. The participant indicates their response in each trial by pressing a left or right arrow button located below the row of arrows on the iPad screen. The software generates a raw score based on a combination of accuracy and reaction time, which is then converted to a standard score.

##### 3. NIH Toolbox Dimensional Change Card Sort Test

The dimensional change card sort test measures cognitive flexibility. Participants view a target image in the center of the iPad screen. Below the target image are two response images: one matches the color of the target image and the other matches the shape of the target image. Before the images are presented, the word “SHAPE” or “COLOR” is displayed on the screen. If SHAPE precedes the images, participants are to press the response image that matches the shape of the target image. If COLOR precedes the images, participants are to press the response image that matches the color of the target image. There are 30 trials, 23% of which are color trials. The software generates a raw score based on a combination of accuracy and reaction time, which is then converted to a standard score.

#### Episodic Memory

##### 1. NIH Toolbox Picture Sequence Memory Test (PSMT)

A sequence of 15 images is displayed on the iPad screen. After the sequence finishes, the participant is instructed to recall the sequence of pictures. Participants move images on the screen in the order they remember them being presented. The participant then completes a second trial consisting of 18 images, which includes the same 15 images as the first trial and adds 3 new images in the middle of the sequence. The raw score is the correct number of adjacent pairs the participant places for each trial, which is then converted to a standard score by the software.

##### 2. Wechsler Memory Scale (WMS-IV) – Verbal Paired Associates (VPA) Subtest [28]

The examiner reads a list of 14 word pairs to the participant. Some of the word pairs consist of related words (e.g. sock-shoe), while other pairs do not (e.g. laugh-stand). In the first part of the test (VPA 1), the list of word pairs is read to the participant four times. The word pairs are presented in a different order each time. After each presentation of the list, the examiner says the first word of each pair, and the participant is instructed to verbally respond with the second word of the pair. Participants receive feedback for every response. If a response is incorrect, they are reminded of the correct answer. Participants receive a point for each item they correctly respond to. There are 56 points possible.

The second part of the subtest (VPA 2) occurs approximately 25 minutes after VPA 1. This is a surprise memory test. In VPA 2-Recall, the examiner says the first word of each pair, and the participant is instructed to respond with the second word of the pair. Participants do not receive feedback. Their raw score is the number of items they correctly respond to out of 14 items.

In VPA 2-Recognition, the examiner reads a list of 40 word pairs. The participant is instructed to indicate if the stated word pair is one of the pairs they were presented with earlier. Their raw score is the number of items they correctly respond to out of 40 items.

##### 3. Rivermead Behavioral Memory Test (RBMT) – Story Subtest [29]

In Part 1 (immediate recall), participants are verbally presented with a brief story consisting of 5 sentences. They are instructed to listen carefully to the story and then tell the examiner as much of the story they can remember. Part 2 (delayed recall) is a surprise memory test that occurs approximately 20 minutes after Part 1. Here, participants are again asked to tell the examiner as much of the story they can remember. Their raw score is the number of “ideas” they correctly remember out of 21 possible ideas.

#### Crystallized Intelligence

##### 1. NIH Toolbox Picture Vocabulary Test (PV)

This test provides a measure of general vocabulary knowledge. The test utilizes Computer Adaptive Testing, in which each question is dependent on the participant’s response to the previous question. Participants hear an audio recording of a word and four pictures are displayed on the iPad screen. They are instructed to select the picture that best matches the meaning of the word they heard. The software generates a raw score using Item Response Theory, which is then converted to a standard score.

##### 2. NIH Toolbox Oral Reading Recognition Test (OR)

This is a measure of reading ability. A word is presented on the iPad screen, and the participant is instructed to read the word out-loud. Using a pronunciation guide, the examiner scores the response as correct or incorrect. Like the PV test, this test utilizes Computer Adaptive Testing. The software generates a raw score using Item Response Theory, which is then converted to a standard score.

### fMRI Session Protocol

Functional MRI data is collected using a 3T General Electric Discovery Magnetic Resonance System with an 8-channel head coil at the University of Michigan’s Functional MRI Laboratory. The two fMRI sessions, each lasting approximately 45 minutes, include the acquisition of a structural image and task-based and resting state functional data. The first task the participants complete is the somatosensory task. The somatosensory task is always presented first so that the devices delivering the stimulation can be removed after this task. The auditory task is always presented second so that the task preceding the resting state scan (the third scan) is the same across all participants. The visual and motor tasks are the last two tasks, and the order of these tasks is counterbalanced across participants.

T2*-weighted images for all four functional tasks are collected using a 2D Gradient Echo pulse sequence with the following parameters: Repetition Time (TR) = 2000 ms; Echo Time (TE) = 30 ms; flip angle = 90°; Field of View (FOV) = 220 × 220 mm; 180 volumes; 43 axial slices with thickness = 3 mm and no spacing.

The specific sequence of scans is described below.

#### (1) 3-Plane Localizer

The localizer is generated with a 2D Gradient Echo pulse sequence with FOV = 320 × 320 mm and slice thickness = 10 mm with no spacing; acquisition time = 30 seconds.

#### (2) T1-weighted Overlay

The overlay is generated with a 2D T1-weighted Fluid-Attenuated Inversion Recovery (FLAIR) pulse sequence with the following parameters: TR = 3173.1 ms; TE = 24.0 ms; Inversion Time (TI) = 896 ms; flip angle = 111°; FOV = 220 × 220 mm; 43 axial slices with thickness = 3 mm and no spacing; acquisition time = 100 seconds.

#### (3) Somatosensory Task

The vibrotactile somatosensory task lasts six minutes and uses two Cortical Metrics Brain Gauge Pro MRI-compatible tactile stimulators (one for each hand), controlled using inhouse Microsoft Visual Studio scripts. The task consists of six 20-second blocks of right index and middle finger stimulation, six 20-second blocks of left index and middle finger stimulation, and twelve 10-second blocks of no stimulation. Each stimulation block consists of twenty 500 ms vibrations interleaved with 500 ms of no vibration to create a pulsing sensation. Each vibration block is followed by a no-stimulation block. The vibration blocks are pseudorandomized, and the block order is the same for all participants. A fixation cross is presented on the screen for the duration of the task. Target trials consist of the 500 ms vibration delivered to one finger instead of both fingers, and there are 6 target trials during the task (3 for each hand). The participant is instructed to press a button with their right thumb every time a target trial occurs (a Current Designs 2-button fiber optic response unit is attached to the right-hand stimulator to collect response data). A target trial occurs approximately once every minute, and there is never more than one target trial in a given block.

#### (4) Auditory Task

The auditory task lasts 6 minutes and consists of six 20-second blocks of foreign speech clips, six 20-second blocks of instrumental music clips; and twelve 10-second blocks of no sound. Each speech block consists of a 20-second segment of a news reporter speaking in a foreign language. The languages used are Creole, Macedonian, Marathi, Persian, Ukranian, and Swahili. Only one language is used per block, and participants are screened to ensure they are unfamiliar with the languages used. Each music block consists of a 20-second segment of instrumental music. Each speech and music block is followed by a no-sound block. The speech and music blocks are pseudorandomized, and the block order is the same for all participants. A fixation cross is presented on the screen for the duration of the task. Target trials consist of a beep interjected into the speech or music, and there are 6 target trials presented throughout the task – 3 for speech and 3 for music. Participants are instructed to press a button with their right index finger every time they hear a target trial. There is a target trial approximately once every minute, and there is never more than one target trial in a given block. Sound is presented through an Avotec Conformal Headset, and responses are collected via a Celeritas 5-button fiber optic response unit. Heart rate is collected via a pulse oximeter placed on the left middle finger.

#### (5) Resting State

T2*-weighted functional resting state data is collected with a 2D Gradient Echo pulse sequence with the following parameters: TR = 2000 ms; TE = 30 ms; flip angle = 90°; FOV = 220 × 220 mm; 240 volumes; 43 axial slices with thickness = 3 mm and no spacing. The resting state acquisition time is 8 minutes. Participants are instructed to relax, keep their eyes open and focus on a fixation cross presented for the duration of the scan. Heart rate is collected via a pulse oximeter placed on the left middle finger.

#### (6) Visual Task

The visual task lasts for 6 minutes and consists of six 20-second blocks of images of male faces, six 20-second blocks of images of houses, and twelve 10-second blocks of a fixation cross. Each block consists of the stimulus presented for 500 ms with an interstimulus interval (ISI) of 500 ms. Every face and house block is followed by a fixation block. The face and house blocks are pseudorandomized, and the block order is the same for all participants. Target trials are images of female faces for face blocks, and images of apartment buildings for house blocks. Participants are instructed to press a button with their right index finger every time they see a target trial. There are 6 target trials presented throughout the task – 3 for the face blocks and 3 for the house blocks. A target trial is presented approximately once every minute, and there is never more than one target trial in a given block. Responses are collected via a Celeritas 5-button fiber optic response unit. Heart rate is collected via a pulse oximeter placed on the left middle finger.

#### (7) Motor Task

The motor task lasts six minutes and consists of six 20-second blocks of a left-pointing arrow, six 20-second blocks of a right-pointing arrow, and twelve 10-second blocks of a fixation cross. Each block consists of the stimulus presented for 500 ms with a 500 ms ISI. Every arrow block is followed by a fixation block. The arrow blocks are pseudorandomized, and the block order is the same for all participants. Participants are instructed to press a button with their right thumb every time they see a right-pointing arrow and to press a button with their left thumb every time they see a left-pointing arrow. Unlike the visual, auditory and somatosensory tasks, the motor task does not contain target trials, since participants are already making active responses. Responses are collected via a Celeritas 5-button fiber optic response unit. Heart rate is collected via a pulse oximeter placed on the left middle finger.

#### (8) High-resolution Structural Image

A high-resolution T1-weighted structural image is collected using a 3D fast spoiled gradient echo (SPGR) BRAVO pulse sequence with the following parameters: TR = 12.2 ms; TE = 5.2 ms; TI = 500 ms; flip angle = 15°; FOV = 256 × 256 mm; 156 axial slices with thickness = 1 mm and no spacing; acquisition time = 5 minutes.

### MRS/DWI Session Protocol

Magnetic resonance spectroscopy (MRS) and Diffusion Weighted Imaging (DWI) data are collected on a different day using the same MRI scanner and head coil as the fMRI scanning session. This session lasts approximately 1.5 hours and consists of the following sequence of scans:

#### (1) 3-Plane Localizer

The localizer is collected using the same parameters as in the fMRI session.

#### (2) T1-weighted Structural Image

The structural image is collected using the same parameters as in the fMRI session.

#### (3) Diffusion Weighted Image (DWI)

DWI data are collected using a diffusion-weighted 2D dual spin echo pulse sequence with the following parameters: TR = 7250 ms; TE = 2.5 ms; FOV = 240 × 240 mm; 32 diffusion directions; 60 axial slices with thickness = 2.4 mm and 0.1 mm spacing. Acquisition time is approximately 10 minutes.

#### (4) Magnetic Resonance Spectroscopy

We collect GABA edited MR spectra from six cortical voxels using a MEGA-PRESS sequence [30] with the following parameters: TR = 1800 ms; TE = 68 ms (TE1 = 15 ms, TE2 = 53 ms); 256 transients (128 ON interleaved with 128 OFF) of 4,096 data points; spectral width = 5 kHz; frequency selective editing pulses (14 ms) applied at 1.9 ppm (ON) and 7.46 ppm (OFF); FOV = 240 × 240 mm; voxel size = 30 × 30 × 30 mm. Acquisition time for each voxel is approximately 8.5 minutes. Voxels are placed in order to maximize overlap with fMRI activation from the corresponding task in the same participant in their own native space. The placements are therefore unique to each participant. To determine voxel placements, we conduct a general linear model (GLM) on each fMRI task, contrasting each condition against rest. For example, in the visual task we compute contrast maps for house vs. fixation and for face vs. fixation. Using these two contrast maps and the T1 structural image, we place the ventral visual voxels to capture the areas of the highest activation (highest beta value) in the house and face areas for each hemisphere. For the auditory voxels we use the contrast maps for speech versus no sound and music versus no sound. For the sensorimotor voxels, we place the left hemisphere voxel to capture activations from the right hand motor task and the right hand somatosensory task. We place the right hemisphere voxel using the left hand activations.

## Drug Study

The goal of the drug study is to explore whether GABA plays a role in age-related neural dedifferentiation. To do so, we manipulate GABA activity pharmacologically using lorazepam (a benzodiazepine) and investigate the effect on neural distinctiveness assessed with fMRI. Lorazepam is an allosteric modulator of the GABA receptor, potentiating its inhibitory function. We hypothesize that increasing GABA activity experimentally will increase neural distinctiveness.

## Methods and design

### Participants

Participants in the drug study do not participate in the main study. All participants are healthy right-handed, native English speakers aged 18-29 (young adults) or 65 and older (older adults). In addition to the major exclusion criteria listed in Table 1, participants are excluded if they have glaucoma, breathing problems, or an allergy to benzodiazepines. They are also excluded if they are undergoing chemotherapy, or have an immune system disorder, or kidney or liver disease. These additional exclusions are enforced due to potential interactions with lorazepam. All sessions take place at the University of Michigan’s Functional MRI Laboratory at the Bonisteel Interdisciplinary Research Building in Ann Arbor, Michigan. Participants are being recruited from the Ann Arbor community and the surrounding area.

### Power Calculations

Tso, Fang, Phan, Welsh, and Taylor [31] is one of the few studies to investigate the effects of lorazepam on blood-oxygen-level dependent imaging (BOLD) responses in healthy adults. They found drug-placebo differences with an approximate effect size of d = 1.15 using a 0.01 mg/kg intravenous dose of lorazepam. To achieve 80% power to detect an effect of this size, a sample of 21 subjects per group would be required. We are therefore recruiting 25 younger adults and 25 older adults for this study.

### Session Design

After completing a telephone screening process and eligibility is confirmed, subjects participate in two separate fMRI sessions. In one of the fMRI sessions, participants are given a placebo pill approximately 1 hour before the scanning session. In the other fMRI session, a 0.5 mg oral dose of lorazepam is administered approximately 1 hour before the session. We determined the lorazepam dosage from a pilot study in which we assessed drowsiness (using a visual analog scale and psychomotor vigilance task) in healthy adults at doses of 0.5 mg, 1 mg, and 2 mg of lorazepam. Participants displayed significant drowsiness at 1 and 2 mg, so we decided to use 0.5 mg for our drug study.

Each fMRI session lasts approximately 45 minutes and includes four different tasks to elicit activation in the visual, auditory, and somatosensory cortices. These fMRI sessions follow the exact same protocol described for the main study. In addition, participants complete a visual analog scale and psychomotor vigilance task just before and immediately following the fMRI scans to assess potential drowsiness. Participants are randomly assigned to one of four session orders as depicted in Table 3. These session orders are used to counterbalance the lorazepam administration and the presentation of the motor and visual fMRI tasks. The method of counterbalancing the fMRI tasks is the same as in the main study.

**Table 3.** Drug pilot session orders

| Group | Session 1 | Session 2 |
| --- | --- | --- |
| 1A | Placebo fMRI<br>fMRI task order: visual-motor | Lorazepam fMRI<br>fMRI task order: motor-visual |
| 1B | Placebo fMRI<br>fMRI task order: motor-visual | Lorazepam fMRI<br>fMRI task order: visual-motor |
| 2A | Lorazepam fMRI<br>fMRI task order: visual-motor | Placebo fMRI<br>fMRI task order: motor-visual |
| 2B | Lorazepam fMRI<br>fMRI task order: motor-visual | Placebo fMRI<br>fMRI task order: visual-motor |

### fMRI Session Protocol

Image acquisition parameters for the fMRI session are the same as in the main study. Subjects participate in two fMRI sessions on separate days: one with placebo and one with lorazepam.

## Preprocessing and analysis pipelines

### Magnetic Resonance Imaging

#### Anatomical MRI

We use surface-based methods as implemented in FreeSurfer to construct a cortical surface for each participant from their high-resolution T1-weighted anatomical image.

#### Functional MRI

##### Task-based fMRI data preprocessing

FreeSurfer and FSFAST are used to perform the preprocessing and first-level analyses of the fMRI data [32]. Preprocessing procedures include motion correction, and spatial smoothing using a Gaussian kernel with full width half maximum (FWHM) of 5mm.

##### Multi-voxel pattern analysis (MVPA)

Neural distinctiveness is assessed using MVPA in functionally defined regions of interest (ROIs). Neural responses are first estimated using a GLM implemented in FSFAST. For each task, responses to the two experimental conditions (visual task: faces and houses; auditory task: speech and music; motor task: left and right finger tapping; tactile task: left and right vibrotactile stimulation) are modelled using a block design, with models including separate regressors for each of the experimental blocks convolved with a canonical hemodynamic response function.

Using FreeSurfer’s Cortical Parcellation technique, four bilateral anatomical masks, one for each task are created for each participant using their T1-weighted structural image. Parcellation results are reviewed and, if necessary, manually corrected. For the visual task, this mask includes the fusiform gyrus and the parahippocampal gyrus. For the auditory task, it includes superior temporal gyrus, transverse temporal gyrus, bank of the superior temporal sulcus, and supramarginal gyrus. And for the motor and tactile tasks this mask includes the precentral gyrus, postcentral gyrus and supramarginal gyrus. Next, in-house MATLAB code is used to combine information from the anatomical masks and the task GLMs to create a functional ROI for each task and participant. First, vertices within each participant’s anatomical mask are sorted based on activation level for one of the experimental conditions vs. rest. Then, a second list is created by sorting vertices within the anatomical ROI based on their activation level for the other experimental condition vs. rest. Finally, the functional ROI is defined by alternating between the two sorted lists, adding the most active voxel (that has not already been included) for the first experimental condition, then adding the most active voxel that has not already been included for the other condition, and so on. This procedure continues until the target functional ROI size is reached. This approach was chosen in order to include voxels activated by both conditions, without biasing the ROI to have more voxels associated with either condition.

Next, the activation estimates within each participant’s functional ROI are used to measure the distinctiveness of multi-voxel representations for conditions of interest in each experimental task. Inspired by Haxby and colleagues [33], we compute Pearson correlations to estimate how similar activation patterns to the same stimulus type are within the functional ROI (e.g., how similar are activation patterns evoked by different face blocks? How similar are activation patterns evoked by different house blocks?). We then average all of the within-condition correlations to get an estimate of within-condition reliability. We also compute correlations between activation patterns evoked by different conditions (e.g., how similar is a face-evoked pattern to a house-evoked pattern?) and average all of the between-condition correlations to get an estimate of between-condition similarity. Finally, we define neural distinctiveness as the difference between the average within-condition correlation and the average between-condition correlation. This neural distinctiveness measure is the difference between two correlations and can therefore range from −2 (very low neural distinctiveness, indicating that between-condition correlations are actually higher than within-condition correlations) to +2 (very high neural distinctiveness, indicating that within-condition correlations are much higher than between-condition correlations). This approach is used rather than alternative classification methods (i.e., support vector machines) that only produce a few distinct accuracy values and that are prone to ceiling effects. Neural distinctiveness is measured separately for each task and each participant.

##### Resting-state fMRI data preprocessing

Preprocessing of the resting-state fMRI data is performed using SPM12 [34]. Preprocessing steps include slice-time correction, realignment, segmentation of structural images, normalization into standard MNI space, and spatial smoothing. The Artifact Detection Toolbox (ART) [35] is used to account for head motion in the scanner. An image is flagged as an outlier if 1) head displacement in the x, y, or z direction is greater than 0.5 mm from the previous frame; 2) the rotational displacement is greater than .02 radians from the previous frame; or 3) the global mean intensity of an image is greater than 3 standard deviations from the mean image intensity of the entire scan. Outliers are included as nuisance covariates in the first-level GLM.

Additional denoising on the resting-state data is performed using the SPM compatible CONN toolbox [36]. Data are first filtered using a temporal band-pass filter of .008 to .09 Hz to ensure analyses focus on the frequency band of interest and higher frequency sources of noise are excluded. For additional noise reduction, the anatomical component-based correction method, aCompCor, is used. This method models the influence of noise as a voxel-specific linear combination of multiple empirically estimated noise sources by producing principal components from noise ROIs and subsequently including them as nuisance parameters in the first level GLM. To do this, each participant’s structural image is segmented into white matter (WM), grey matter (GM), and cerebrospinal fluid masks (CSF). Next, the WM and CSF masks are eroded by one voxel to minimize partial voluming with GM. These eroded WM and CSF masks are thereafter used as noise ROIs.

The signals from all ROIs are extracted from the unsmoothed functional images to avoid potential “spillage” of the BOLD signal from nearby regions. Residual head motion parameters (three rotations, three translations and six parameters representing their first-order temporal derivatives) and signals from WM and CSF are regressed out during the calculation of functional connectivity maps.

### Magnetic Resonance Spectroscopy

#### Preprocessing and analysis

For each of the six MRS voxels collected, GABA concentrations are quantified using the MEGA-PRESS difference spectra using Gannet 3.0 [37], which is specifically targeted for GABA edited MRS. Time domain data are frequency- and phase-corrected using spectral registration and filtered with a 3-Hz exponential line broadening and zero-filled by a factor of 16. Gannet models the GABA peak using a five-parameter Gaussian model between 2.19 and 3.55 μm, and the water peak using a Gaussian-Lorentzian function. In all analyses, metabolite concentration values are scaled to water, and expressed in institutional units (IU). GABA estimates are then corrected for the fraction of the voxel that is cerebrospinal fluid (CSF) and white matter (WM) as opposed to grey matter (GM) (using SPM12 segmentation), and for the different water relaxation times in CSF, WM, and GM as described in Harris et al. [38].

### Diffusion Weighted Images

#### Preprocessing

Diffusion-weighted imaged (DWI) are preprocessed using MRtrix [39]. Preprocessing includes Echo-planar imaging (EPI) correction, motion correction, and bias field correction. DWIs are intensity normalized across subjects based on the median b = 0 s/mm^2^ intensity within a white matter mask [40]. Images are up-sampled to an isotropic voxel size of 1.3 mm using b-spline interpolation [40]. Fiber orientation distributions (FODs) are computed using robust constrained spherical deconvolution (rCSD) [41]. A group average response is used to estimate FODs in all subjects, as described in Raffelt et al. [40]. Registration is performed using FODs at I_max_ 4, 100 equally distributed apodised point spread functions during FOD reorientation, with displacement field smoothing (Gaussian kernel σ^2^ = 1), velocity field smoothing (Gaussian kernel σ^2^ = 3), and an initial gradient step of 0.2. A white matter template analysis fixel mask was generated with an fmls peak value of 0.15. Whole brain probabilistic tractography is then performed on the FOD template generating 20 million streamlines and SIFT is applied with an output of 2 million streamlines.

#### Fixel-based analysis

We are performing fixel based analyses (FBA) of fiber density (FD), fiber bundle cross-section (FC), and fiber density and cross section (FDC). Measures of FD, FC and FDC are computed as described in Raffelt et al. [42]. For the FC and FDC analyses, we include intra-cranial volume (computed from T1-weighted images, using FreeSurfer) as a nuisance covariate. We are also performing Connectivity-based Fixel Enhancement (CFE) using 2 million streamlines and default parameters (smoothing = 10 mm FWHM, C = 0.5, E = 2, H = 3; taken from Raffelt et al. [43]). In this analysis, C is a constant weighting how structurally connected fixels (hypothesized to share underlying axons) contribute to the enhancement of other fixels. The H parameter allows increasing weight to an extent (i.e., a group of connected fixels) at higher test-statistic thresholds, and E influences how much the extent influences the enhancement as it scales in size. For further details of these parameters, see Raffelt et al. [43].

## Discussion

Tens of millions of otherwise healthy people are already experiencing age-related behavioral impairments, and based on population projections, that number is going to grow significantly in the coming years. However, there are substantial individual differences in the degree of cognitive decline that people experience as they age. The Michigan Neural Distinctiveness (MiND) project investigates the underlying causes of age-related impairments, the consequences of these impairments, and what distinguishes people who age gracefully from those who don’t. Developing an understanding of the source of individual differences in aging is an important step in designing interventions that could slow, or conceivably even reverse, the behavioral impairments associated with healthy aging. This study is innovative in at least four ways.

First, we are collecting fMRI, MRS, and behavioral measures in the same subjects, a rare combination. Doing so will allow us to directly investigate the relationship between neural distinctiveness (assessed using fMRI), GABA levels (assessed using MRS), and age-related behavioral declines (assessed using psychophysical and assessment techniques).

Second, we are using multivariate pattern analysis (MVPA) techniques that allow us to study neural activation *patterns* rather than treating every brain voxel as independent (as more traditional univariate techniques do). Multivoxel analyses often reveal information that univariate analyses miss [44,45].

Third, we are explicitly investigating *individual differences* in neurochemistry, neural representation, and behavioral performance. A lot of work on the neuroscience of aging investigates group differences between young and older subjects and ignores individual differences between subjects in the same age group. However, some older subjects experience significantly greater behavioral declines than others, and these individual differences could provide important insights into the underlying causes of age-related behavioral declines.

Finally, the proposed research offers the potential to inspire a new approach to therapy for age-related behavioral impairments. The majority of current interventions focus on behavioral training (either cognitive training or physical exercise) [46–49]. However, if we can demonstrate that reductions in GABA levels play an important role, then biological interventions (e.g., GABA agonists and related pharmaceuticals) might also be a fruitful therapeutic path to pursue.

## List of abbreviations

ART: Artifact Detection Toolbox
BIN: Buildings in Noise
BOLD: Blood-oxygen-level dependent imaging
CFE: Connectivity-based Fixel Enhancement
CSF: Cerebrospinal fluid
dB: Decibel
DWI: Diffusion weighted imaging
EPI: Echo-planar imaging
FBA: Fixel based analyses
FC: Fiber bundle cross-section
FD: Fiber density
FDC: Fiber density cross-section
FIN: Faces in Noise
FLAIR: Fluid-Attenuated Inversion Recovery
fMRI: Functional magnetic resonance imaging
FODs: Fiber orientation distributions
FOV: Field of view
FWHM: Full width half maximum
GABA: Gamma-aminobutyric acid
GLM: General linear model
GM: Grey matter
ISI: Interstimulus interval
IU: Institutional units
MiND: Michigan Neural Distinctiveness
MoCA: Montreal Cognitive Assessment
MRS: Magnetic resonance spectroscopy
MVPA: Multi-voxel pattern analysis
OIN: Objects in Noise
OR: Oral Reading
PSMT: Picture Sequence Memory Test
PTA: Pure tone average
PV: Picture Vocabulary
RBMT: Rivermead Behavioral Memory Test
rCSD: Robust constrained spherical deconvolution
ROI: Region of interest
ScIN: Scenes in Noise
SNR: Signal-to-noise ratio
SPGR: Spoiled gradient echo
TE: Echo time
TI: Inversion time
TR: Repetition time
VPA: Verbal Paired Associates
WAIS: Wechsler Adult Intelligence Scale
WIN: Words in Noise
WM: White matter
WMS: Wechsler Memory Scale

## Declarations

### Ethics approval and consent to participate

The MiND project is approved by the University of Michigan Medical School Institutional Review Board (IRBMED; Study ID HUM00103117).

### Consent for publication

Not applicable.

### Availability of data and material

Data sharing is not applicable to this article as no datasets were generated or analyzed during the current study.

### Authors’ contributions

TP is the PI of the study and received the grant funding the project.

HG and MS were the main authors of the manuscript.

HG and EF were major contributors to the coordination of the project and data collection.

MS, BF, and MP were the main contributors to MRS design and analyses.

HG, MS, KC, JC, PL, DP, RDS, and DHW contributed to task design and data analyses.

SFT contributed to drug-related components of the study.

SK contributed to DWI analyses.

All authors read and approved the final manuscript.

## Acknowledgements

Not applicable.

## References

1. Leventhal A, Wang Y, Pu M, Zhou Y, Ma Y. GABA and its agonists improved visual cortical function in senescent monkeys. SCIENCE. 2003 May 2;300(5620):812–5.

2. Carp J, Park J, Polk TA, Park DC. Age differences in neural distinctiveness revealed by multi-voxel pattern analysis. NeuroImage. 2011 May 15;56(2):736–43.

3. Park DC, Polk TA, Park R, Minear M, Savage A, Smith MR. Aging reduces neural specialization in ventral visual cortex. Proc Natl Acad Sci U S A. 2004 Aug 31;101(35):13091–5.

4. Park J, Carp J, Hebrank A, Park DC, Polk TA. Neural specificity predicts fluid processing ability in older adults. J Neurosci. 2010 Jul 7;30(27):9253–9.

5. David-Jürgens M, Churs L, Berkefeld T, Zepka RF, Dinse HR. Differential effects of aging on fore–and hindpaw maps of rat somatosensory cortex. Harris J, editor. PLoS ONE. 2008 Oct 14;3(10):e3399.

6. Spengler F, Godde B, Dinse HR. Effects of ageing on topographic organization of somatosensory cortex. Neuroreport. 1995 Feb 15;6(3):469–73.

7. Villers-Sidani E de, Alzghoul L, Zhou X, Simpson KL, Lin RCS, Merzenich MM. Recovery of functional and structural age-related changes in the rat primary auditory cortex with operant training. Proc Natl Acad Sci. 2010 Aug 3;107(31):13900–5.

8. Payer D, Marshuetz C, Sutton B, Hebrank A, Welsh RC, Park DC. Decreased neural specialization in old adults on a working memory task. Neuroreport. 2006 Apr 3;17(5):487–91.

9. Carp J, Park J, Hebrank A, Park DC, Polk TA. Age-related neural dedifferentiation in the motor system. Wenderoth N, editor. PLoS ONE. 2011 Dec 22;6(12):e29411.

10. Park DC, Lautenschlager G, Hedden T, Davidson NS, Smith AD, Smith PK. Models of visuospatial and verbal memory across the adult life span. Psychol Aging. 2002 Jun;17(2):299–320.

11. Gershon RC, Wagster MV, Hendrie HC, Fox NA, Cook KF, Nowinski CJ. NIH toolbox for assessment of neurological and behavioral function. Neurology. 2013 Mar 12;80(11 Suppl 3):S2–6.

12. NIH Toolbox Scoring and Interpretation Guide for the iPad (2016). NIH Toolbox and PROMIS iPad apps. https://nihtoolbox.desk.com/customer/portal/articles/2437205-nih-toolbox-scoring-and-interpretation-guide. Accessed 30 October 2018.

13. Broadbent DE, Cooper PF, FitzGerald P, Parkes KR. The Cognitive Failures Questionnaire (CFQ) and its correlates. Br J Clin Psychol. 1982 Feb;21(1):1–16.

14. Rast P, Zimprich D, Van Boxtel M, Jolles J. Factor Structure and Measurement Invariance of the Cognitive Failures Questionnaire Across the Adult Life Span. Assessment. 2009 Jun;16(2):145–58.

15. Brainard DH. The Psychophysics Toolbox. Spat Vis. 1997;10(4):433–6.

16. Kleiner M, Brainard DH, Pelli D, Ingling A, Murray R, Broussard C. What’s new in Psychtoolbox-3. Vol. 36. 2007. 1 p.

17. The Mathworks, Inc. MATLAB. Natick, Massachusetts, United States: The MathWorks, Inc.; 2017b.

18. Gold J, Bennett PJ, Sekuler AB. Signal but not noise changes with perceptual learning. Nature. 1999 Nov 11;402(6758):176–8.

19. Brady TF, Konkle T, Alvarez GA, Oliva A. Visual long-term memory has a massive storage capacity for object details. Proc Natl Acad Sci U S A. 2008 Sep 23;105(38):14325–9.

20. Zhou B, Lapedriza A, Khosla A, Oliva A, Torralba A. Places: A 10 Million Image Database for Scene Recognition. IEEE Trans Pattern Anal Mach Intell. 2018 Jun 1;40(6):1452–64.

21. Smits C, Kapteyn TS, Houtgast T. Development and validation of an automatic speech-in-noise screening test by telephone. Int J Audiol. 2004 Jan;43(1):15–28.

22. Holden JK, Nguyen RH, Francisco EM, Zhang Z, Dennis RG, Tommerdahl M. A novel device for the study of somatosensory information processing. J Neurosci Methods. 2012 Mar 15;204(2):215–20.

23. Zhang Z, Francisco EM, Holden JK, Dennis RG, Tommerdahl M. Somatosensory information processing in the aging population. Front Aging Neurosci. 2011;3:18.

24. Nguyen RH, Forshey TM, Holden JK, Francisco EM, Kirsch B, Favorov O, et al. Vibrotactile discriminative capacity is impacted in a digit-specific manner with concurrent unattended hand stimulation. Exp Brain Res. 2014 Nov;232(11):3601–12.

25. Carey LM, Nankervis J, LeBlanc S, Harvey LA. A new functional Tactual Object Recognition Test (fTORT) for stroke clients: Normative standards and discriminative validity. 14th Int Congr World Fed Occup Ther. 2006;

26. Nasreddine ZS, Phillips NA, Bédirian V, Charbonneau S, Whitehead V, Collin I, et al. The Montreal Cognitive Assessment, MoCA: a brief screening tool for mild cognitive impairment. J Am Geriatr Soc. 2005 Apr;53(4):695–9.

27. Wechsler D. Wechsler Adult Intelligence Scale-Fourth Edition. Fourth. San Antonio, TX: Pearson; 2008.

28. Wechsler D. Wechsler Memory Scale -(WMS–IV) technical and interpretive manual. Fourth. San Antonio, TX: Pearson; 2009.

29. Wilson B, Greenfield E, Clare L, Baddeley A, Cockburn J, Watson P, et al. The Rivermead Behavioral Memory Test-Third Edition (RBMT-3). Third. London, UK: Pearson; 2008.

30. Mullins PG, McGonigle DJ, O’Gorman RL, Puts NAJ, Vidyasagar R, Evans CJ, et al. Current practice in the use of MEGA-PRESS spectroscopy for the detection of GABA. NeuroImage. 2014 Feb 1;86:43–52.

31. Tso IF, Fang Y, Phan KL, Welsh RC, Taylor SF. Abnormal GABAergic function and face processing in schizophrenia: A pharmacologic-fMRI study. Schizophr Res. 2015 Oct;168(1–2):338–44.

32. FreeSurfer. http://surfer.nmr.mgh.harvard.edu/. Accessed 30 October 2018.

33. Haxby JV, Gobbini MI, Furey ML, Ishai A, Schouten JL, Pietrini P. Distributed and Overlapping Representations of Faces and Objects in Ventral Temporal Cortex. Science. 2001 Sep 28;293(5539):2425–30.

34. SPM. https://www.fil.ion.ucl.ac.uk/spm/. Accessed 30 October 2018.

35. Artifact Detection Tools (ART). NITRC: NeuroImaging Tools and Resources Collaboratory. https://www.nitrc.org/projects/artifact_detect/. Accessed 30 October 2018.

36. CONN: functional connectivity toolbox. NITRC: NeuroImaging Tools and Resources Collaboratory. https://www.nitrc.org/projects/conn. Accessed 30 October 2018.

37. Gannet [Internet]. [cited 2018 Oct 30]. Available from: http://www.gabamrs.com/

38. Harris AD, Puts NAJ, Barker PB, Edden RAE. Spectral-editing measurements of GABA in the human brain with and without macromolecule suppression: Relationship of GABA+ and MM-suppressed GABA. Magn Reson Med. 2015 Dec;74(6):1523–9.

39. MRtrix [Internet]. [cited 2018 Oct 30]. Available from: https://mrtrix.readthedocs.io/en/latest/index.html

40. Raffelt D, Tournier J-D, Rose S, Ridgway GR, Henderson R, Crozier S, et al. Apparent Fibre Density: A novel measure for the analysis of diffusion-weighted magnetic resonance images. NeuroImage. 2012 Feb;59(4):3976–94.

41. Tournier J-D, Calamante F, Connelly A. Determination of the appropriate *b* value and number of gradient directions for high-angular-resolution diffusion-weighted imaging: APPROPRIATE *b* VALUE AND NUMBER OF GRADIENT DIRECTIONS FOR HARDI. NMR Biomed. 2013 Dec;26(12):1775–86.

42. Raffelt DA, Tournier J-D, Smith RE, Vaughan DN, Jackson G, Ridgway GR, et al. Investigating white matter fibre density and morphology using fixel-based analysis. NeuroImage. 2017 Jan;144:58–73.

43. Raffelt DA, Smith RE, Ridgway GR, Tournier J-D, Vaughan DN, Rose S, et al. Connectivity-based fixel enhancement: Whole-brain statistical analysis of diffusion MRI measures in the presence of crossing fibres. NeuroImage. 2015 Aug;117:40–55.

44. Haynes J-D, Rees G. Decoding mental states from brain activity in humans. Nat Rev Neurosci. 2006 Jul;7(7):523–34.

45. Norman KA, Polyn SM, Detre GJ, Haxby JV. Beyond mind-reading: multi-voxel pattern analysis of fMRI data. Trends Cogn Sci. 2006 Sep;10(9):424–30.

46. Ball K, Berch DB, Helmers KF, Jobe JB, Leveck MD, Marsiske M, et al. Effects of cognitive training interventions with older adults: A randomized controlled trial. JAMA. 2002 Nov 13;288(18):2271.

47. Belleville S, Gilbert B, Fontaine F, Gagnon L, Ménard E, Gauthier S. Improvement of episodic memory in persons with mild cognitive impairment and healthy older adults: evidence from a cognitive intervention program. Dement Geriatr Cogn Disord. 2006;22(5–6):486–99.

48. Colcombe S, Kramer AF. Fitness effects on the cognitive function of older adults: a meta-analytic study. Psychol Sci. 2003 Mar;14(2):125–30.

49. Lautenschlager NT, Cox KL, Flicker L, Foster JK, van Bockxmeer FM, Xiao J, et al. Effect of physical activity on cognitive function in older adults at risk for Alzheimer disease: A randomized trial. JAMA. 2008 Sep 3;300(9):1027.

